# Wound-initiated hair regeneration by adhesive and shrinkable materials

**DOI:** 10.1101/2024.08.27.610012

**Authors:** Shoichiro Kokabu, Kunikazu Tsuji, Ayako Washio, Kazumasa Murata, Mitsushiro Nakatomi, Yusuke Ono, Osamu Kaminuma, Takuma Matsubara

**Affiliations:** Division of Molecular Signaling and Biochemistry, Kyushu Dental University, Kitakyushu, Fukuoka, 803-8580, Japan; (S.K.); (T. M.); Department of Orthopedic Surgery, Tokyo Medical and Dental University, Bunkyo, Tokyo, 113-8519, Japan; (K. T.); Division of Endodontics and Restorative Dentistry, Kyushu Dental University, Kitakyushu, Fukuoka, 803-8580, Japan; (A. W.); (K. M.); Department of Human, Information and Life Sciences, School of Health Sciences, University of Occupational and Environmental Health, Kitakyushu, Fukuoka, 807-8555, Japan; (M. N.); Department of Muscle Development and Regeneration, Institute of Molecular Embryology and Genetics (IMEG), Kumamoto University, Kumamoto, 860-0811, Japan; (Y. O.); Tokyo Metropolitan Institute for Geriatrics and Gerontology, Itabashi, Tokyo, 173-0015, Japan; (Y. O.); Department of Disease Model, Research Institute of Radiation Biology and Medicine, Hiroshima University, Hiroshima, 734-8551, Japan. (O. K.)

## Abstract

Although there is a global demand for hair regrowth, particularly among middle-aged and older individuals, an effective hair growth technology has not yet been established^1^. Hair follicle neogenesis is restricted to the embryonic period, but hair regeneration accompanied by wound healing has been observed under some conditions^2–4^; however, the underlying mechanisms are unclear. Herein, we demonstrated that creating a wound without dermal defects effectively induced postneonatal hair follicle neogenesis. Separating the epidermis from the dermis by topical application of adhesive and shrinkable materials to mouse skin promoted epidermal regeneration, followed by new hair follicle formation. Hair follicle regeneration, accompanied by the upregulation of related genes, can be induced in mice, including middle-aged and aged mice, regardless of species, sex, skin location, or age. The cycle of the regenerated hair eventually synchronized with that of the surrounding physiological hairs. Our new hair regeneration technique based on reproduction of epidermis–dermis interactions provides a novel means to treat hair loss, including androgenetic alopecia.

## Introduction

Globally, there is a strong desire for techniques for hair regrowth among males and females. The global market for alopecia therapy was $7.6 billion in 2020 and is expected to reach $13 billion by 2028^1^. Since natural hair follicle formation is known to occur only during the embryonic period, several attempts to treat alopecia by transplanting hair stem cells and hair follicle organoids have been reported^5–15^. However, there have been reports of postneonatal follicular formation associated with the wound-healing process in rabbits^16^, mice^17^, and humans^18^. A mouse model for investigating wound-induced hair neogenesis (WIHN) has been developed^2^. However, the detailed mechanisms underlying the initiation of WIHN have not been clarified.

An individual hair is generated in a hair follicle through a single hair cycle^19^. The hair follicle, once formed in the embryonic period, repeats the hair cycle, which consists of the catagen, telogen, and anagen phases, throughout its lifetime. During the development of a hair follicle, the epithelium plunges into the mesenchyme, forming a tubular structure^20^. Epithelial–mesenchymal interactions are essential for initiation, and dermal-mesenchymal interactions predominate during the progression of hair follicle formation^21^. No hair follicles formed from chimeric skin that combined the epidermis of the dorsal skin and the dermis of the plantar area. However, hair follicles formed where the dermis of the dorsal skin and the epidermis of the plantar area were combined^22^, suggesting that the mesenchyme-derived dermis plays a key role in achieving postneonatal hair regeneration.

We hypothesized that hair follicle neogenesis can be induced under conditions in which the dermis remains intact during the skin regeneration process. Successful skin wound induction without dermal defects by a newly developed procedure involving the application of adhesive and shrinkable materials to the skin promoted hair regeneration accompanied by de novo hair follicle neogenesis at almost 100% probability, even in middle-aged and aged mice. The regenerating hair cycle eventually synchronizes with the surrounding hair cycle.

## Results

### Shrink material induces hair regeneration

The dorsal hairs of 5-week-old C57BL/6 mice are known to be in the anagen phase. After the hairs were shaved, the skin color became black because the hairs remained in the hair roots in the skin (Fig. 1a). In contrast, the hairs of 8-week-old mice were in the telogen phase and could be easily removed from their roots by shaving. Therefore, the skin revealed its natural skin color after shaving. After application to the shaved dorsal skin in the telogen phase, 10% pyroxylin solution adhered to the skin and then shrank upon drying. Approximately 2 days after application, a skin wound developed at the site of pyroxylin application. Surprisingly, visible hair growth was observed at the wound site approximately 14 days after application. On approximately day 19, hair growth that was essentially at the same level as that of the original, unshaved hairs in the surrounding skin area, both in terms of length and density, was observed (Fig. 1a).

**Fig. 1.**
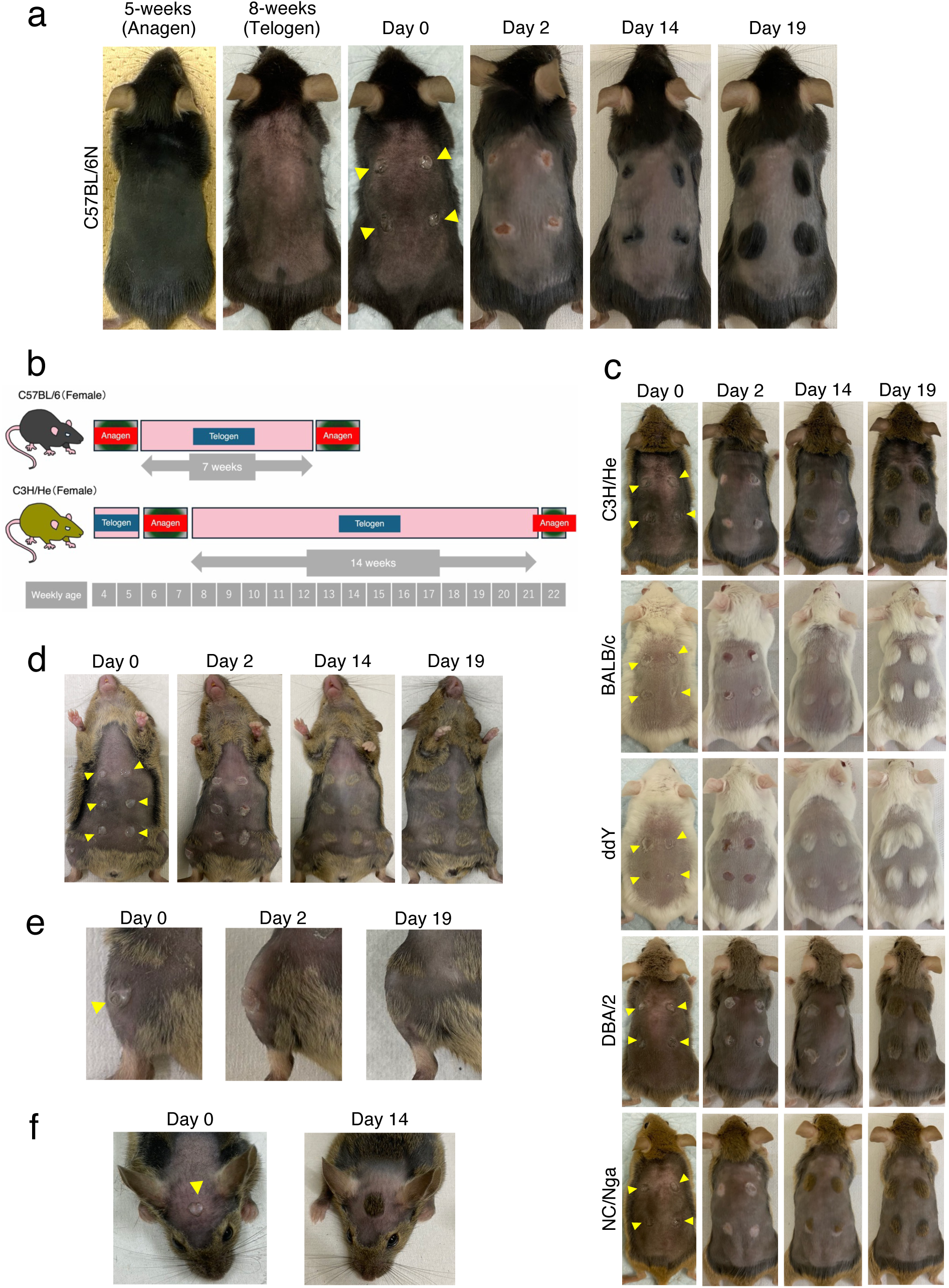
Pyroxylin induces skin wounds followed by hair regeneration. (a) Ten percent pyroxylin solution was applied to the dorsal skin of 5– and 8-week-old C57BL/6N female mice after shaving (yellow arrowhead). Representative photos of 5-week-old mice on day 2 and 8-week-old mice on days 0, 2, 14 and 19 after shaving are shown. (b) The schematics represent the hair cycle of C57BL/6N or C3H/He female mice, as previously reported. (c) Pyroxylin was applied to the dorsal skin of 8-week-old C3H/He, BALB/c, ddY, DBA/2, and NC/Nga mice after shaving (yellow arrowhead). Representative photos on days 0, 2, 14, and 19 are shown. (d-f) Pyroxylin was applied to the skin of the abdomen (d), lower leg (e), and head (f) after shaving (yellow arrowhead). Representative photos on days 0, 2, 14, and/or 19 are shown. (a-f) All mice examined exhibited the same trend (n=6).

The hair cycle, including the length and timing of telogen and anagen, varies between the sexes and in mouse species^23,24^. In addition to C57BL/6, C3H/He has been used for wound healing and hair growth research^4,23,24^. The hair cycles of these two mouse strains are different (Fig. 1b; http://www.jslc.co.jp/pdf/mouse/2020/019_C3H_HeNSlc). However, there was no significant difference in the level or time course of hair regeneration following pyroxylin application between the C57BL/6N and C3H/He mouse strains (Fig. 1a and c) or between male and female mice (data not shown).

The abdominal and limb hair characteristics significantly differ from those of the dorsal hair. The skin of the head and back develops from distinct germ layers during embryonic development^25,26^. Therefore, pyroxylin-induced hair regeneration was examined in these regions of C3H/He mice, which are characterized by a long telogen period and a tightly synchronized hair cycle^23^ (Fig. 1b). After the development of wounds by pyroxylin application to the abdomen (Fig. 1d), lower limb (Fig. 1e), and head (Fig. 1f) after shaving, hairs regenerated in all the regions with almost the same time course as that observed in the dorsal skin. However, in addition to the lack of beard regeneration, de novo hair growth was not observed in hairless regions, such as the plantar area, following pyroxylin application (data not shown).

The dose-dependent ability of pyroxylin to induce hair regeneration was examined. Pyroxylin at concentrations greater than 1.25% induced essentially the same level of hair regeneration, although obvious wound and hair regeneration was not observed at concentrations less than 0.625% (Fig. 2a). The effects of adhesion– and shrinkage-related materials other than pyroxylin were examined. The application of bisphenol A-glycidyl methacrylate (Bis-GMA)-based composite resins with low-or high-shrinkage characteristics, polymethyl methacrylate (PMMA), solvent-based styrene-butadiene rubber, and cyanoacrylates created wounds and induced hair regeneration in the dorsal skin, similar to those observed with pyroxylin treatment. Water-soluble materials such as ethylene-vinyl acetate copolymer emulsions and polyvinyl alcohol aqueous solutions can also induce hair regeneration. However, adaptation of pure liquid components, including MMA + 4-META or MMA alone, did not induce hair regeneration (Fig. 2b). To eliminate the involvement of pure chemical irritation in hair regeneration, we applied high concentrations of acids, such as salicylic acid^27^, trichloroacetic acid^28^, glycolic acid^29^, and lactic acid^30^, which are frequently used as chemical peeling reagents, to the skin. No apparent wound or hair regeneration was induced by any type or concentration of acid examined (Fig. 2c).

**Fig. 2.**
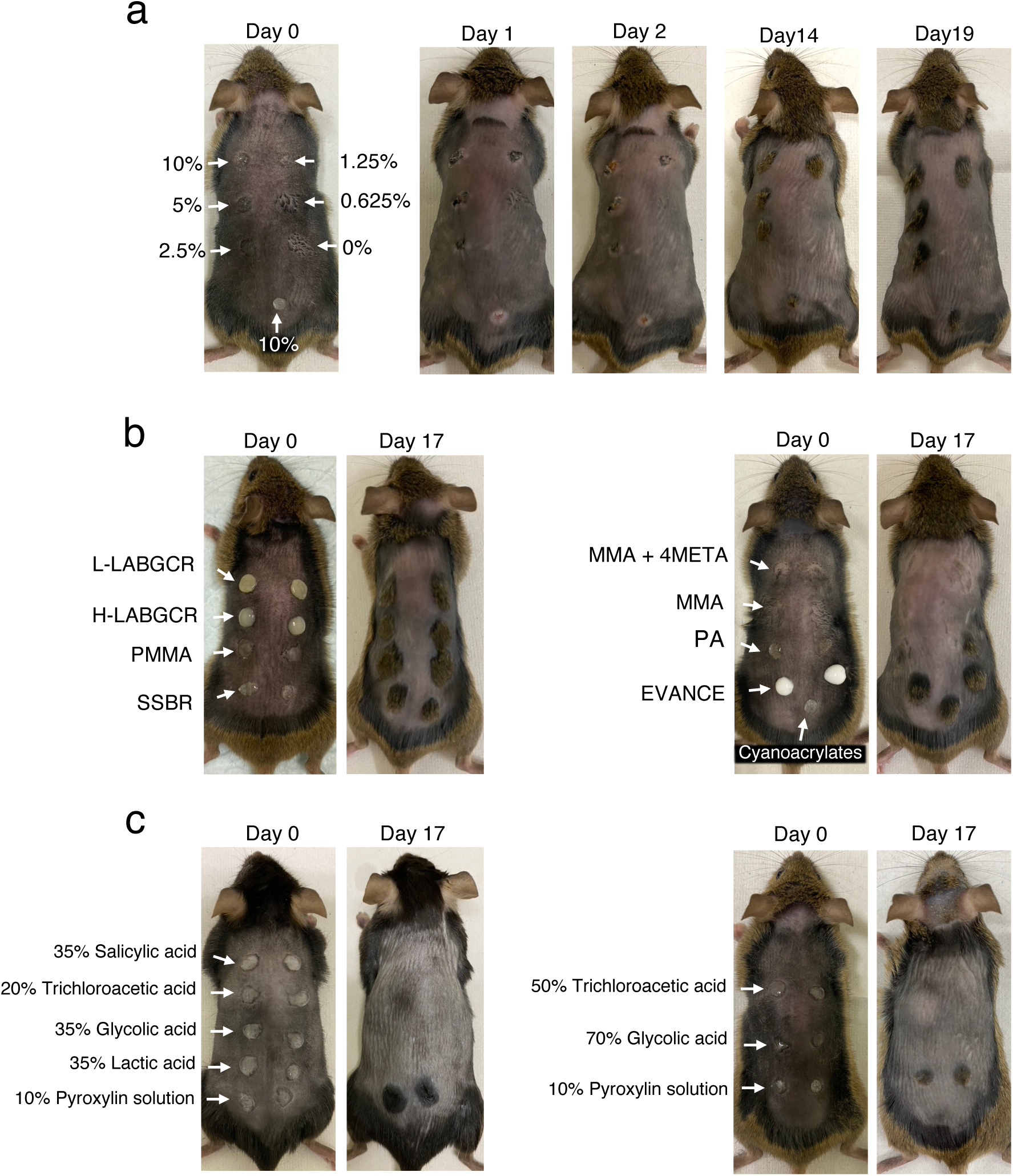
Hair regeneration is induced in the skin by the application of adhesive and shrinkable materials. (a) Zero (ethanol: diethyl ether = 1:1 weight), 0.625, 1.25, 2.5, 5, or 10% pyroxylin solution was applied (yellow arrowhead) to the dorsal skin of 8-week-old C3H/He mice after shaving. Representative photos on days 0, 2, 14, and 19 are shown. (b and c) Low-or high-shrinkage light-activated Bis-GMA-based composite resins (L-LABGCR or H-LABGCR), PMMA, SSBR, MMA, MMA + 4-META, PA, EVACE, and cyanoacrylates (b). After shaving, 35% salicylic acid solution, 20% or 50% trichloroacetic acid solution, 35% or 70% glycolic acid solution, and 35% lactic acid solution (c) were applied to the dorsal skin. Representative photos of three independent experiments are shown.

We next performed histological analysis during pyroxylin-induced wound generation and hair regeneration in the dorsal skin. Three hours after pyroxylin treatment, peeling of the epidermis from the dermis was restricted to the area where pyroxylin was applied (Fig. 3a and b). One day later, the epidermis became necrotic, accompanied by an accumulation of inflammatory cells. On day 2, in addition to the beginning of epidermal regeneration with necrotic old epidermis detachment, invasion of part of the regenerated epidermis into the dermis was observed. On day 5, the invasion of the epidermis toward the deeper dermis and the generation of new hair follicles and sebaceous glands were detected. On day 9, hair shafts were identified in further-grown new hair follicles (Fig. 3c). Hair placodes, which are located on dermal condensates and represent the primordia of hair follicles, can be identified by alkaline phosphatase (ALP) activity^31^. The hair follicle primordium was indicated in the tips of the newly invaded epidermis by increased ALP activity (Fig. 3d). The expression of hair follicle formation– and neogenesis-related genes such as Sox9^32^, Lhx2^32^, Bmp7^33^, Wnt10b^2^, Lef1^2^, Sonic hedgehog (Shh)^2^ and Gli1^33^ gradually increased in the dorsal skin following pyroxylin application, peaked at 9 to 14 days, and then decreased (Fig. 3e). These data suggested that separating the epidermis from the dermis is crucial for activating hair follicle formation– and neogenesis-related genes and resulting in the progression of hair regeneration. We named this novel hair follicle neogenesis process epidermis separation-induced hair neogenesis (ESHN).

**Fig. 3.**
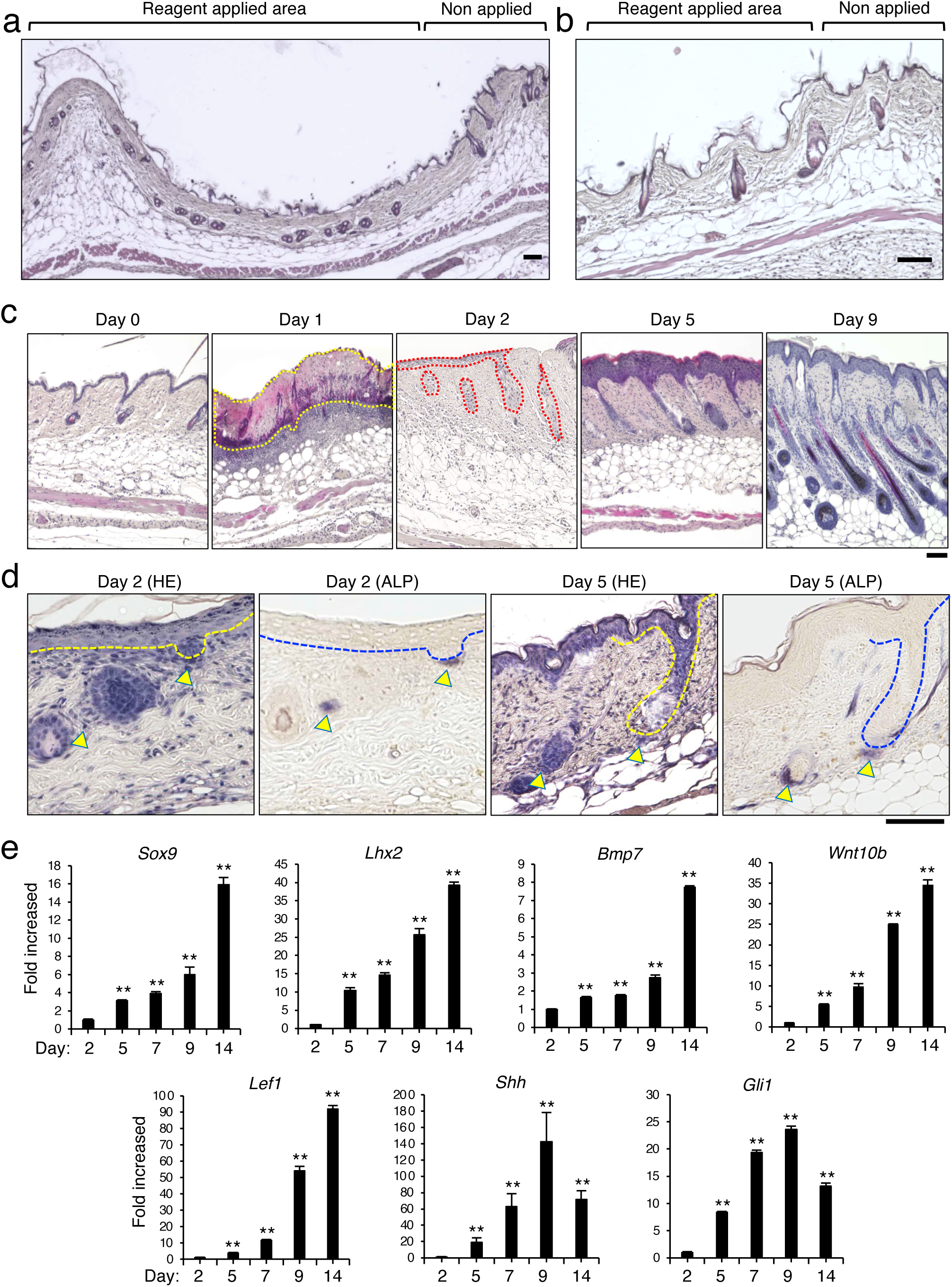
Hair follicle neogenesis following the separation of the epidermis and dermis. (a and b) Representative H&E staining images of wounds on dorsal back skin 3 hours after pyroxylin application (a) and a corresponding magnified image (b) are shown. (c) Representative H&E staining images of dorsal back skin wounds on days 0, 1, 2, 5, and 9 are shown. The yellow and red dotted lines indicate the necrotic epidermis and growing panicles, respectively. (d) Representative H&E staining (left) and ALP activity staining (right) images of wounds on the dorsal back skin on days 2 or 5 are shown. The left and right images are consecutive specimens. The arrowhead indicates the ALP activity-positive area. The yellow and blue dotted lines indicate growing panicles and hair follicles, respectively. Representative images (a-d) with the same trend from 6 individual mice are shown. (a-d) Scale bars indicate 100 μm. (e) mRNA levels of Sox9, Lhx2, Bmp7, Wnt10b, Lef1, Shh, and Gli1 in the dorsal back skin of wounds on 2, 5, 7, 9, or 14 were determined by real-time PCR. The data are expressed as the mean ± SEM of triplicate measurements. \*\**p* < 0.01, compared with the day 2 sample. Essentially, the same results were obtained from three independent mice.

### ESHN occurs in middle-aged and aged mice

The demand for hair regenerative medicine is increasing in the middle-aged and aged life stages because hair loss is an age-related process^34^. Based on the correlation between mouse and human life spans, 56– and 78-week-old C57BL/6J mice are regarded as middle-aged and aged, respectively^35^. The dorsal hair cycles of 56– and 78-week-old C57BL/6J mice are in the 4th telogen phase^24^. Therefore, we next examined ESHN in these mice. Pyroxylin-induced hair regeneration was observed in 56-week-old mice over the same time course as that observed in 8-week-old mice (Figs. 1a, c and 4a). Interestingly, similar to the surrounding original hairs, the regenerated hairs were a mixture of black and gray hairs (Fig. 4b). Hair follicle neogenesis was detected by histological examination (Fig. 4c). ESHN occurred even in 78-week-old mice (Fig. 4d and e).

**Fig. 4.**
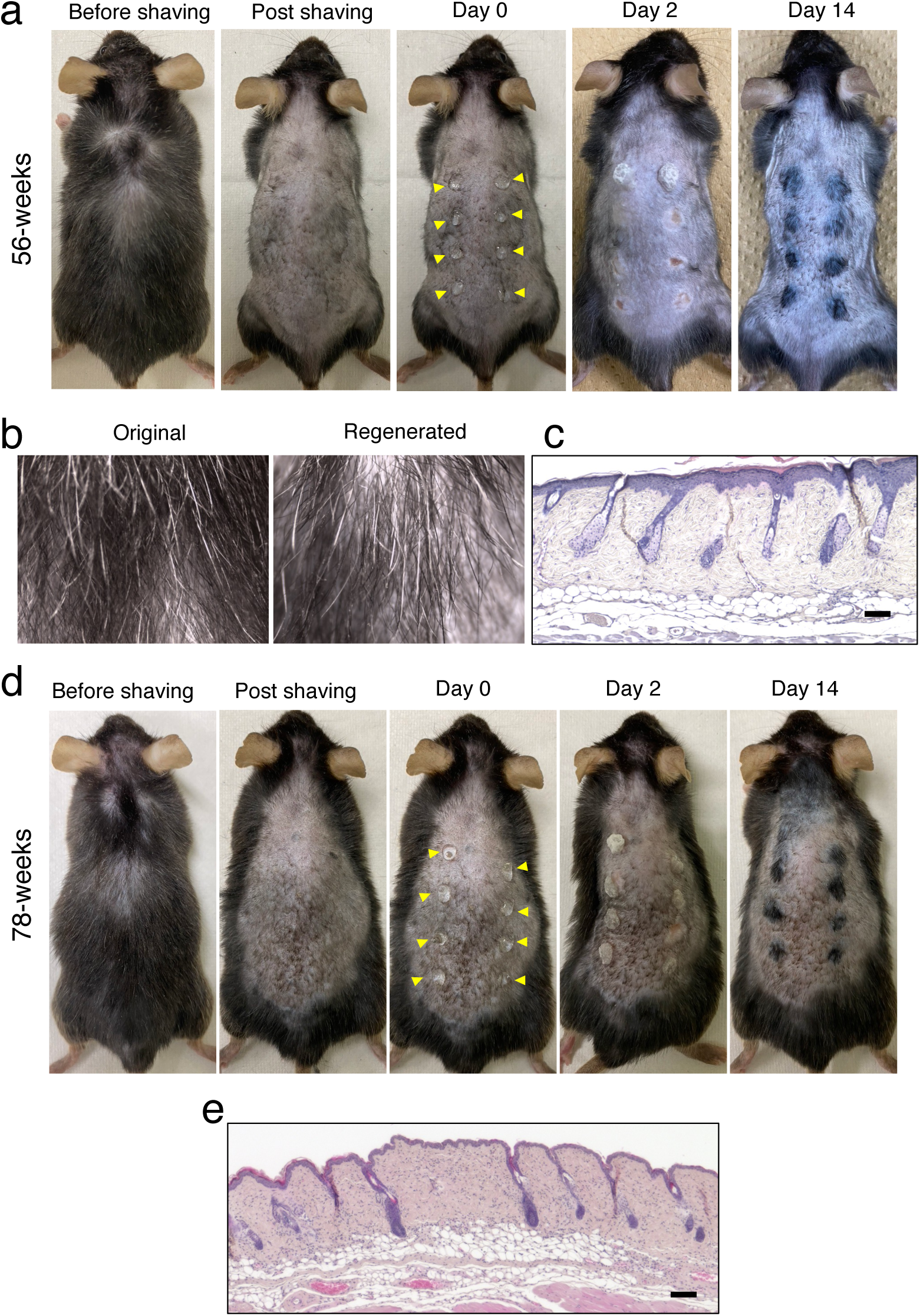
ESHN in middle-aged and aged mice. (a-e) Ten percent pyroxylin solution was applied (yellow arrowhead) to the dorsal hair of middle-aged (a-c) and aged (d, e) C57BL/6J male mice after shaving. (a, d) Representative photos on days 0, 2, and 14 are shown. (b, d) Representative photos of original resided hairs (left) and regenerated hairs (right: on day 20) observed under stereomicroscopy are shown. (c, e) Representative H&E staining images of dorsal back skin wounds on day 5 are shown. Scale bars indicate 100 μm. (a-e) All mice examined exhibited the same trend (n=6).

### ESHN synchronizes surrounding hair cycle

We further examined the cycle of hair regeneration following repeated shaving. After the length of the regenerated hairs reached approximately the same level as that of the surrounding hairs on day 19, the regenerated hairs were shaved daily. This process enabled the determination of the hair cycle based on the skin color. The black color of the hair regeneration area after shaving on day 19 gradually faded with each subsequent shaving. By the 25th day, the color became almost indistinguishable from the surrounding skin color (Fig. 5a). Corresponding to the skin color alteration, long hair follicles in an anagen phase with high density were observed in the dorsal skin on day 19. Following repeated daily shaving, the length and density of the follicles tended to decrease, suggesting a transition to the catagen phase. After 7 repeated shavings on day 25, the follicles almost completely entered the telogen phase (Fig. 5b). The hair regeneration process following repeated pyroxylin treatment was examined. Pyroxylin was reapplied to the area where ESHN once occurred during the telogen phase. Hair regeneration was induced in the second pyroxylin application area at the same level as that in the first application area. Essentially, the same level of hair regeneration was also evoked by the third pyroxylin application (Fig. 5c). When ESHN was induced in 4-week-old C3H/He mice that were in the first physiological telogen phase, we observed hair growth two weeks prior to the beginning of the first physiological anagen phase. Following the start of the first physiological anagen phase at approximately day 17, the hair also began to grow from the shaved area without ESHN induction. Based on the skin color after daily shaving from days 20 to 32, the hair cycle in the area of ESHN induction seemed to enter the telogen phase earlier than that in the area retaining the physiological hair cycle. However, the difference mostly disappeared at approximately day 90. Thus, the cycle of ESHN-induced hairs eventually synchronized with that of the surrounding physiological hairs during the second anagen phase (Fig. 5d).

**Fig. 5.**
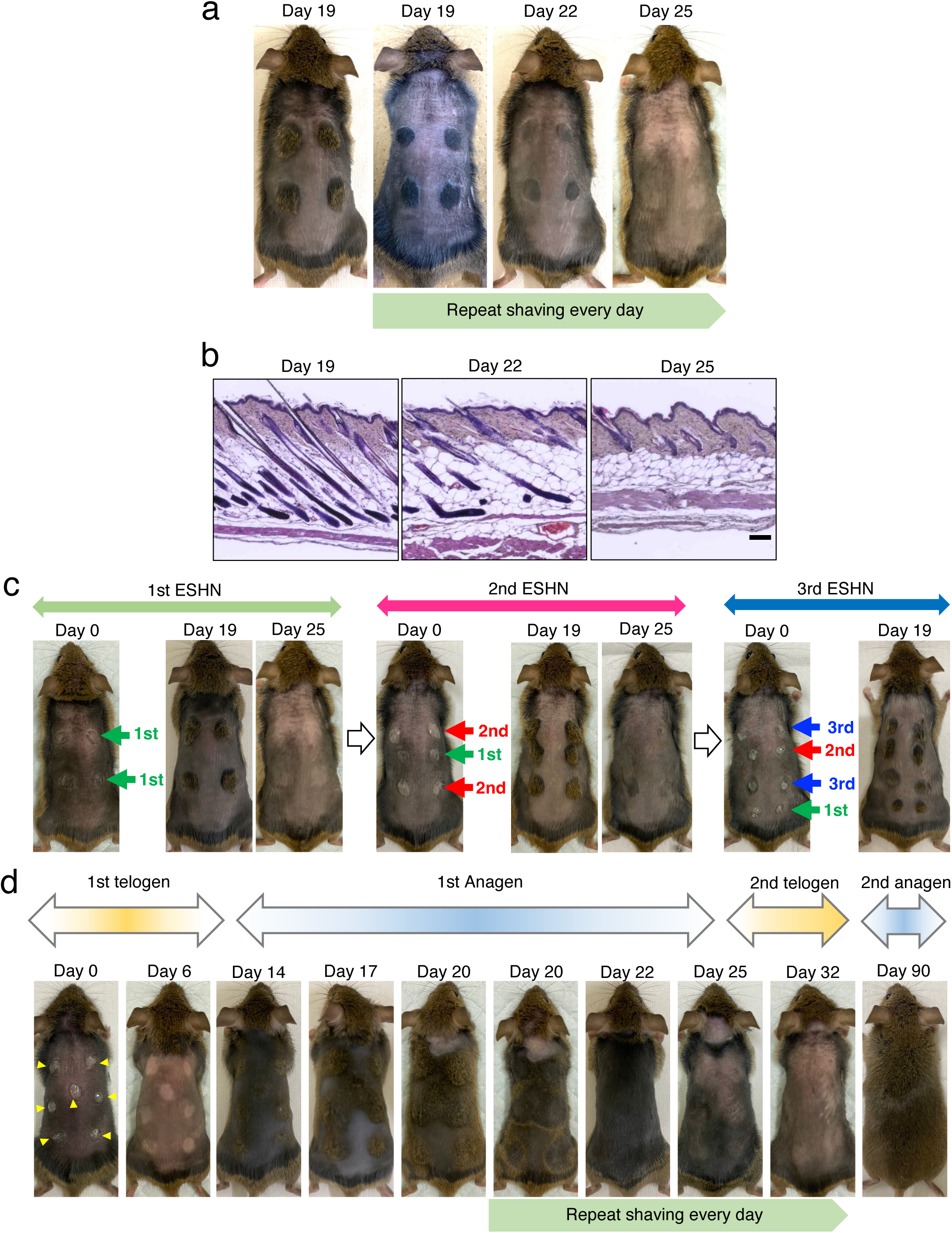
Hair cycle synchronization in regenerated and surrounding hairs. (a) Ten percent pyroxylin solution was applied to the dorsal hair of 8-week-old C3H/He mice after shaving. Beginning on day 19, the regenerated hairs were shaved every day. Representative photos on days 19, 22, and 25 are shown. (b) Representative H&E staining images of dorsal back skin wounds on days 19, 22, and 25 are shown. (c) Ten percent pyroxylin solution was applied to the dorsal hair of 8-week-old C3H/He mice after shaving (green arrow). The photos are shown on day 0, 19, or 25 (1st ESHN). A 10% pyroxylin solution was also applied at the same location for the first time (2nd time; red arrow). The photos are shown on day 0, 19, or 25 (2nd ESHN). Again, a 10% pyroxylin solution was applied to the same place for the 2nd time (3rd time; blue arrow). The photos are shown on day 0, 19, or 25 (3rd ESHN). (d) Pyroxylin was applied to the dorsal hair of 4-week-old C3H/He mice after shaving (yellow arrowhead). Beginning on day 20, the regenerated hair was shaved every day for 2 weeks. Representative photos on days 0, 6, 14, 17, 20, 22, 25, 32, and 90 are shown. All mice examined exhibited the same trend (n=6).

## Discussion

In this study, we developed a new technique for inducing hair follicle neogenesis in postneonatal mice. The simple application of adhesive and shrinkable materials to the skin to create a wound without dermal damage initiated the regeneration of hairs with a normal hair cycle even in middle-aged and aged mice.

Based on previous observations showing hair follicle formation during the wound-healing process in the skin of several species^16–18^, Ito et al. developed a mouse model of WIHN^2^. When a full-thickness excision of a certain size was created on the dorsal skin of mice in the first or second telogen phase, hair follicle neogenesis occurred in the center of the wound healing area within 2 weeks, and visible hair growth was observed after approximately 4 weeks^2^. The contribution of Wnt signaling to WIHN was further elucidated. Several procedures for follicular regeneration without creating wounds have also been reported. Gat et al. reported that mice expressing stabilized β-catenin controlled by an epidermal promoter undergo a process resembling de novo hair morphogenesis^36^. Essentially, the same results were reported by Lo Celso et al., although both mentioned tumorigenesis based on the maintained activation of Wnt/β-catenin signaling^36,37^. The activation of Hedgehog signaling in adjacent epithelial and stromal cells induces new hair follicle and tumor formation^38^.

Several signaling cascades responsible for postneonatal hair formation have been suggested from these previous studies. However, the key event triggering hair follicle neogenesis has still not been identified even in the WIHN model. To explore the black box, we focused on epithelial-mesenchymal interactions. The hair follicle, along with teeth, sweat glands, mammary glands, and salivary glands, is classified as an ectodermal organ. These organs are formed via epithelial–mesenchymal interactions. In particular, mesenchymal cells initiate the organ fate-determining process^21,39^. The initial signal arising from the dermis causes the formation of fair follicles during embryogenesis^40^. Therefore, we hypothesized that the mesenchyme-derived dermis, which remains intact during wound healing, retains the information necessary to initiate hair follicle formation. This hypothesis was proven, at least in part, by our present study demonstrating hair follicle formation following the regeneration of epithelium-derived epidermis above the intact dermis. Yang et al. demonstrated hair follicle neogenesis by artificial dermis transplantation after the creation of a full-thickness skin defect of a size that would not normally trigger WIHN^4^, further supporting our results and hypothesis. The ESHN we established in this study might mimic the follicle formation process during embryogenesis.

Our ESHN procedure has several advantages compared to the previous WIHN model^2–4^. Thus, hair regeneration could be induced simultaneously in almost all the mice examined in short periods. ESHN can be induced multiple times in the skin during the telogen phase. ESHN potentially contributes to the development of hair follicle regenerative methods, e.g., in combination with microsurgery or laser procedures by which the epidermis layer alone can be removed. A new lineage tracking system that enables the identification of the origin of new hair follicles may be helpful to achieve this goal. Organ regeneration procedures based on epithelial-mesenchymal interactions may also be applied to generate regenerative medicines for other ectodermal organs, such as teeth and some glands.

For development of hair follicle regenerative medicine, it is also important to understand the hair cycle process in middle-aged and aged skin environments. Like all other organs, age-related alterations occur in the skin. For example, aged skin is characterized by atrophy (thinning), fragility, dyspigmentation and delayed wound healing^41–43^. In addition, senescent skin cells accumulate progressively with age and impact skin structure and function^44^. During the hair cycle, as the anagen phase gradually shortens, many hair follicles that go through the catagen phase tend to remain in the telogen phase due to aging^45^. This alteration is the main cause of age-related thinning and hair loss, including androgenetic alopecia. In the mouse model of WIHN established by Ito et al., hair follicle neogenesis was inducible only in 7-to 8-week-old or younger mice^2^. However, our present findings demonstrated ESHN not only in young mice but also in middle-aged and aged mice. Moreover, ESHN actively elicited the transition of the hair cycle to the anagen phase from the telogen phase. Although further investigation into the detailed molecular mechanisms underlying the active telogen‒ anagen transition, e.g., by employing spatial transcriptome analysis, is needed, the results strongly suggested that the hair cycle, even under aged conditions, is regulated independently of other age-related alterations observed in the skin. The ESHN-based active hair cycle transition system is promising for the development of hair growth drugs with novel mechanisms useful even in aged humans.

## Methods

### Animals

Mice were purchased from Japan SLC, Inc. (Hamamatsu, Japan). The mouse strains used were BALB/c, ddY, DBA/2, NC/Nga, C3H/He, NOD/SCID, and C57BL/6N. We also purchased 56-or 78-week-old C57BL/6J mice from The Jackson Laboratory Japan, Inc. (Yokohama, Japan). All studies were performed in accordance with the guidelines of and approved by the Experimental Animal Care and Use Committee of Kyushu Dental University (approval numbers #22-004, #22-020, #22-022, #23-006, #23-014, and #23-016).

### Adhesive and shrinkable materials

Various concentrations of pyroxylin solution (ethanol: diethyl ether = 1:1 wight) (FUJIFILM Wako Chemicals, Osaka, Japan), hydrosoluble ethylene-vinyl acetate copolymer emulsion (EVACE, ALTEC, Shiga, Japan), polyvinyl alcohol aqueous solution (PA, Yamato Co., Ltd., Tokyo, Japan), cyanoacrylates (Toagosei Company, Limited, Tokyo, Japan), solvent-based styrene butadiene rubber (SSBR, Cemedine Co., Ltd., Tokyo, Japan), light-activated bis-GMA-based composite resin (LABGCR, Nippon Shika Yakuin Co., Ltd., Shimonoseki, Japan), methyl methacrylate (MMA, Shofu, Inc., Kyoto, Japan), 4-methacryloxy ethyl trimelitate anhydride (4-META, Sun Medical Company), and polymethyl methacrylate (PMMA) powder (Sun Medical Company, Ltd., Shiga, Japan) mixed with MMA + 4META were used for dorsal application. PMMA powder was mixed with MMA and 4-META to initiate polymerization immediately before dorsal application. Salicylic acid (FUJIFILM Wako Chemicals), trichloroacetic acid (FUJIFILM Wako Chemicals), glycolic acid (FUJIFILM Wako Chemicals), and lactic acid (FUJIFILM Wako Chemicals) diluted in polyethylene glycol 300 (FUJIFILM Wako Chemicals) were also used.

### Material application to the skin

Mice were anesthetized under general anesthesia using a triad of anesthetics: medetomidine (Nippon Zenyaku Kogyo Co., Ltd., Fukushima, Japan) (0.75 mg/kg), midazolam (Astellas Pharma, Inc., Tokyo, Japan) (4 mg/kg), and butorphanol (Meiji Seika Pharma Co., Pharma, Inc., Tokyo, Japan) (5 mg/kg)^46–48^. The dorsal hair was shaved using clippers, and materials were applied as 7-8-mm-diameter circular sites. Light-activated bis-GMA-based composite resins were polymerized by light-emitting diode irradiation immediately following application.

### Quantitative real-time PCR

Total RNA was isolated from the skin tissues using a FastGeneTM RNA Basic Kit (Nippon Genetics, Tokyo, Japan) and reverse-transcribed into cDNA using a high-capacity cDNA reverse transcription kit (Applied Biosystems, Waltham, MA, USA). SYBR Green-based quantitative polymerase chain reaction (qPCR) was performed using PowerUp SYBR (Thermo Fisher Scientific, Waltham, MA, USA) and the QuantStudio 3 Real-Time PCR System (Thermo Fisher Scientific). Relative quantification was performed by the *Δ*CT method using *Gapdh* and *Tbp* as the housekeeping genes for normalization. Primer information was provided as supramental information.

### Histopathological examination

Skin samples were fixed with 4% paraformaldehyde (Nacalai Tesque, Inc., Kyoto, Japan) in PBS, dehydrated through an ethanol and xylene series, embedded in paraffin, and cut into 4-μm sections^49^. After deparaffinization, hematoxylin (FUJIFILM Wako Chemicals) and eosin (FUJIFILM Wako Chemicals) (H&E) staining or ALP activity staining were performed. ALP staining substrates were purchased and used in accordance with the instruction manual (Sigma‒Aldrich, St. Louis, MO). The sections were imaged with a Keyence BZ-X800 (Keyence, Osaka, Japan). The shafted hairs were observed using a stereomicroscope (Leica EZ4 HD, Leica, Wetzlar, Germany).

### Statistical analysis

qPCR data are expressed as the mean ± standard error of the mean (SEM) for the fold changes in gene expression compared with that of the control mice. The data were analyzed using one-way ANOVA with multiple comparisons. *p* < 0.01 was considered to indicate statistical significance.

## Data availability

All data reported in this paper will be provide by corresponding author.

## Code availability

This paper does not report original code.

## Acknowledgements

The authors thank William N. Addison for comments on the manuscript. This work was supported by Supported by Kitakyushu Foundation for the Advancement of Industry,

Science and Technology (FAIS) 2023 (to S. K.), The YMFG Regional Enterprise Support Foundation 2024 (to S. K.), Suzuken Memorial Foundation 2023 (to S. K.), The Joint Usage/Research Center for Developmental Medicine, IMEG, Kumamoto University (to S. K.).

## Author contributions

S.K., K. T., A. W. and K. M. conceived of and designed the study. S.K., A.W., and K. M. are performed experiments. S. K., K. T., M. N., Y. O., O. K., and M. T. analyzed the data. S. K. and O. K. wrote the manuscript.

## Additional information

Correspondence and requests for materials should be addressed to Shoichiro Kokabu, D.D.S, Ph.D., Division of Molecular Signaling and Biochemistry, Kyushu Dental University, 2-6-1 Kitakyushu, Manazuru, Kitakyushu, Fukuoka 803-8580, Japan, Telephone: +81-93-285-3047, FAX: +81-93-285-6000; Reprints and permissions information is available at www.nature.com/reprints.

